# Genetic interaction analysis reveals that *Cryptococcus neoformans* utilizes multiple acetyl-CoA-generating pathways during infection

**DOI:** 10.1101/2022.03.16.484561

**Authors:** Katy M. Alden, Andrew J. Jezewski, Sarah R. Beattie, David Fox, Damian J. Krysan

## Abstract

*Cryptococcus neoformans* is an important human fungal pathogen for which the external environment is its primary niche. To cause infection and disease, *C. neoformans* must adapt to a plethora of conditions and stresses inherent to the host environment. Of these stresses, the role of central carbon metabolism has been of interest not only at the level of fundamental pathobiology but also as a potential target for new antifungal drug therapy. Previous work has shown that two non-essential acetyl-CoA metabolism enzymes, ATP-citrate lyase (*ACL1*) and acetyl-CoA synthetase (*ACS1*), play important roles in *C. neoformans* infection. Here, we took a genetic interaction approach to studying the interplay between these two enzymes along with an enzyme initially called *ACS2* but which we have found is an acetoacetyl-CoA synthetase; we have renamed the gene 2-**k**eto**b**utyryl **C**oA synthetase 1 (*KBC1*) based on its biochemical activity and the systematic name of its substrate. *ACL1* and *ACS1* represent a synthetic lethal pair of genes based on our genetic interaction studies. Double mutants of *KBC1* with either *ACS1* or *ACL1* do not have significant synthetic phenotypes in vitro, although we find that deletion of any one of these enzymes reduces fitness within macrophages. Importantly, the *acs1*Δ *kbc1*Δ double mutant has significantly reduced fitness in the CNS relative to either single mutant as well as WT (~2 log_10_ CFU reduction in fungal burden), indicating the important role these enzymes play during infection. The expression of both *ACS1* and *KBC1* is increased in vivo relative to in vitro conditions. The *acs1*Δ mutant is hypersusceptible to fluconazole in vivo despite its minimal in vitro phenotypes. These data not only provide insights into the in vivo mechanism of action for a new class of antifungal Acs inhibitors but also into metabolic adaptations of *C. neoformans* to the host environment.

**Author Summary:** The adaptation of environmental fungal pathogens to the mammalian host is critical to pathogenesis. Of these adaptations, the remodeling of carbon metabolism is particularly important. Here, we generated a focused set of double mutants of non-essential genes (*ACL1, ACS1, KBC1*) involved in acetyl-CoA metabolism and examined their fitness in vitro and during CNS infection. From these studies, we found that all three enzymes play important roles during infection but that the role of *ACS1/KBC1* was minimal in vitro. Consistent with these observations, the expression of *ACS1* and *KBC1* was increased in vivo relative to standard in vitro conditions. Furthermore, strains lacking both *ACL1* and *ACS1* were not viable. These data clearly show that *C. neoformans* employs multiple carbon metabolism pathways to adapt to the host environment. They also provide insights into the potential mechanism of action for anti-cryptococcal Acs inhibitors.

## Introduction

*Cryptococcus neoformans* is an environmental yeast and an accidental human pathogen (1). As a causative agent of Cryptococcosis, it is a major cause of disease in immunocompromised patients. Globally, HIV/AIDS remains the dominant risk factor associated with the development of cryptococcal meningoencephalitis, the primary disease manifestation (2). Understanding the underlying mechanisms that allow this environmental yeast to become a pathogen has been a topic of intense interest to the medical mycology field (3). The majority of these studies have focused on the factors or characteristics that appear to be required for virulence: the so-called Big Three of host temperature tolerance, melanin formation and capsule production (4). Based on large-scale, pheno- and genotyping of diverse clinical and environmental isolates, the expression of these virulence traits is, indeed, strongly associated with disease in humans (5). Despite this correlation, significant variation in virulence toward human patients and in mammalian models of cryptococcosis is observed within highly related isolates that express all three of the canonical virulence traits (6). Accordingly, it is now well-recognized that additional infection-related traits remain to be discovered and characterized (7). Recently, we and others have become interested in the hypothesis that a deeper understanding of this variation may emerge by exploring the traits and processes required for *C. neoformans* to transition from an environmental niche to the host (8).

Pioneering transcriptional profiling of *C. neoformans* during infection of the murine lung (9) and rabbit cerebrospinal fluid (10, 11) by Kronstad, Perfect and co-workers have clearly demonstrated that the expression of many genes related to central carbon metabolism is altered relative to in vitro conditions. These labs have gone on to show that specific enzymes involved in glucose metabolism and acetyl-CoA production are critical for either establishment of infection or progression of disease (12, 13). Griffiths et al. demonstrated that ATP-citrate lyase (*ACL1*), an enzyme that converts citrate generated in, and transported from, the mitochondria into cytosolic acetyl-CoA, is required for robust growth in glucose, production of virulence traits, and virulence in the pulmonary mouse model of cryptococcosis (13). In addition, acetyl-CoA synthetase (*ACS1*) was found to be dispensable for growth on glucose but important for replication on non-fermentable carbon sources such as acetate, glycerol, and ethanol (9). Although deletion of *ACS1* reduced virulence in the mammalian model of cryptococcosis (9), it caused delayed mortality relative to the near avirulence of *acl1*Δ mutants (13). Thus, full fitness of *C. neoformans* during mammalian infection appears to require acetyl-CoA be generated from a variety of metabolic sources.

Our interest in acetyl-CoA metabolism in *C. neoformans* was prompted by the discovery of a small molecule inhibitor of Acs1 that showed antifungal activity and, in particular, synergy with fluconazole against *C. neoformans* in vivo (14). Yeast such as *Saccharomyces cerevisiae, Candida albicans* and *Candida glabrata* lack ACL orthologs; instead, they express two copies of ACS, one which is essential (15). As discussed above, *C. neoformans* lacking *ACS1* are viable. Although the viability of the *acs1*Δ mutant is likely due to the presence of *ACL1*, which is present in other fungi such as *Aspergillus* spp., or the presence of additional isoforms of *ACS*. Based on the annotation of Hu et al., *C. neoformans* has two genes related to *ACS1* and, accordingly, these were annotated as *ACS2* and *ACS3* (9). *ACS1* was clearly related to the essential Sc*ACS2* isoform which is localized to the nucleus and cytoplasm (Fig. 1A); *ACS3* was much less related to ACSs and shows sequence similarity to the *S. cerevisiae* oxalyl CoA synthetase *PSC60* based on BLAST search. CnAcs2 was more closely related to CnAcs1, ScAcs1, and ScAcs2 than Cn*ACS3*. A key distinction between CnAcs1 and CnAcs2 is that CnAcs1 has tryptophan in the conserved substrate binding pocket while CnAcs2 has a glycine (Gly 434). Other members of the **A**cyl-CoA/**N**RPS/**L**uciferase (ANL) family with glycine at this position are acetoacetyl-CoA synthetases that convert the ketone body acetoacetate to acetoacetyl-CoA (16). Acetoacetyl-CoA can then be split to two molecules of acetyl-CoA by acetoacetyl-CoA lyases such as *POT1*/CNAG_00490.

**Figure 1:**
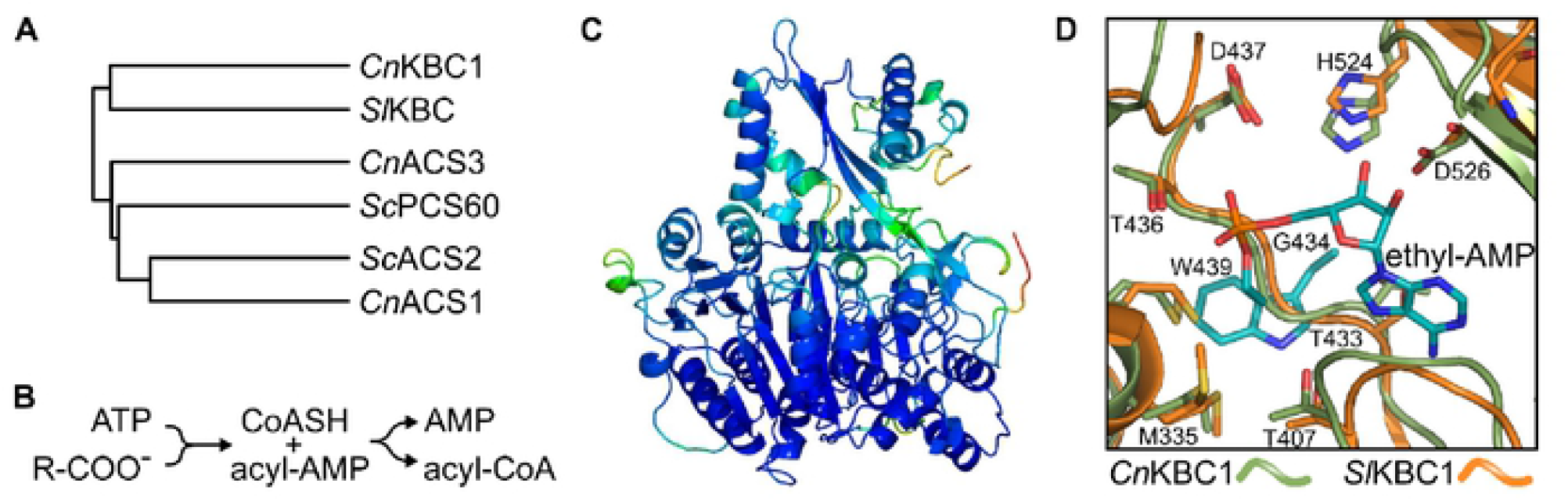
Sequence and structure homology indicate CNAG_02045 encodes a 3-keto-butanoyl-CoA synthetase, *KBC1*. Phylogenetic relationship of ANL-family acyl-CoA synthetases generated using iTOL from a multiple sequence alignment using COBALT (A). Reaction diagram of ANL-family acyl-CoA synthetases (B). AlphaFold2 model of *C. neoformans* Kbc1 generated using the Colab server and colored by spectrum (red-to-blue with increasing level of confidence) of predicted LDDT (local distance difference test) per residue. Image rendered using PyMol v.2.4.OaO (C). Overlay of AlphaFold2 model of *C. neoformans* Kbc1 (green), *S. lividans* Kbc1 (PDB 4WD1, orange), and key components from *C. neoformas* Acs1 (PDB 7KNO, cyan). Ethyl-AMP inhibitor from *Cn*Acs1 shown to demonstrate relative position of the active site. *Cn*Acs1 “Trp wall” residue W439 side-chain occupies same position as the backbone atoms for *Cn*Kbc1 (G434) and S/Kbc1 (G422), effectively closing off the pocket to larger substrates.

Here, we report that CNAG_02045 is an acetoacetyl-CoA synthetase and propose a revised name of 3-**k**eto-**b**utanoyl-**C**oA synthetase (*KBC1*) that is based on the IUPAC systematic chemical name for its substrate and is consistent with current gene naming conventions. We hypothesized that a tertiary route for acetyl-CoA production through *KBC1* may play a role in virulence and that coordinated regulation of *ACS1, ACL1* and *KBC1* is important for the virulence of *Cryptococcus neoformans*. Consistent with that hypothesis, genetic interaction analysis shows that *KBC1* negatively interacts with *ACS1* to reduce fitness of *C. neoformans* during infection of the brain. We also find that loss of *ACS1* and *KBC1* increase susceptibility of *C. neoformans* to fluconazole treatment in vivo to a much greater extent than observed in vitro. During replication in macrophages, loss of any of the three acetyl-CoA related genes leads to reduced fitness, consistent with the low nutrient status of the phagolysosome. We also show that the expression of *KBC1* and *ACS1* is increased in vivo and are coordinately regulated by lipids in vitro. These data support the conclusion *C. neoformans* adapts by utilizing multiple carbon sources to generate the requisite amount of acetyl-CoA for efficient replication.

## Results

### CNAG_02045 is a specific acetoacetyl-CoA synthetase

As discussed above, the sequence of *ACS2*/CNAG_02045 matches much better with an acetoacetyl-CoA synthetase than an *ACS* (Fig. 1A). ANL-family acyl-CoA synthetases (17) catalyze a two-step reaction in which the carboxylic acid is first coupled with ATP to generate an acyl-AMP ester and release pyrophosphate (Fig. 1B). In the second step of the reaction, CoASH condenses with the acyl-AMP to release AMP and the acyl-CoA product. We generated a homology model of CNAG_02045 based on the structure of *Streptomyces lividans* acetoacetyl-CoA synthetase (Fig 1C, reference 16). Using this model, we compared the predicted substrate binding site of CNAG_02045 to the crystal structure of Acs1 bound to an ethyl-AMP substrate mimic (Fig. 1D, reference 18). In Acs1, W439 acts as a “wall” that limits the size of the acetate binding pocket and prevents larger alkyl substituted carboxylic acids from functioning as substrates (17, 18). CNAG_02045 has a Gly (G433) at the position analogous to W439 and, in contrast, the backbone of the peptide occupies the space where the acetyl group would normally bind. As such the predicted substrate binding site of CNAG_02045 is structurally quite distinct from Acs1. No structures of acetoacetyl-CoA synthetases with substrates or substrate mimics have been reported. Mitchell et al. have proposed that a highly conserved threonine (T436) may function as an H-bond donor to the aceoacetate substrate; however, that hypothesis has not been experimentally tested (16).

A 6X-histidine-tagged CNAG_02045 construct was expressed in *E. coli* and the protein purified by IMAC (Fig. S1A). We used a coupled, continuous assay based on detection of pyrophosphate-release that we previously optimized for CnAcs1 (18) to assay CNAG_02045 for acetoacetyl-CoA synthetase activity (Fig. S1B). We observed robust activity (Fig. 2A) that was dependent on both enzyme concentration (Fig. S1C) and the presence of all reaction substrates (Fig. S1D). The K_m_ values for the substrates were: acetoacetate (239± 97 μM); ATP (40±4 μM); and CoASH (55±11 μM) (Fig. 2B). We also observed significant substrate inhibition for CoASH (K_i_ 1130 ± 210 μM).

**Figure 2:**
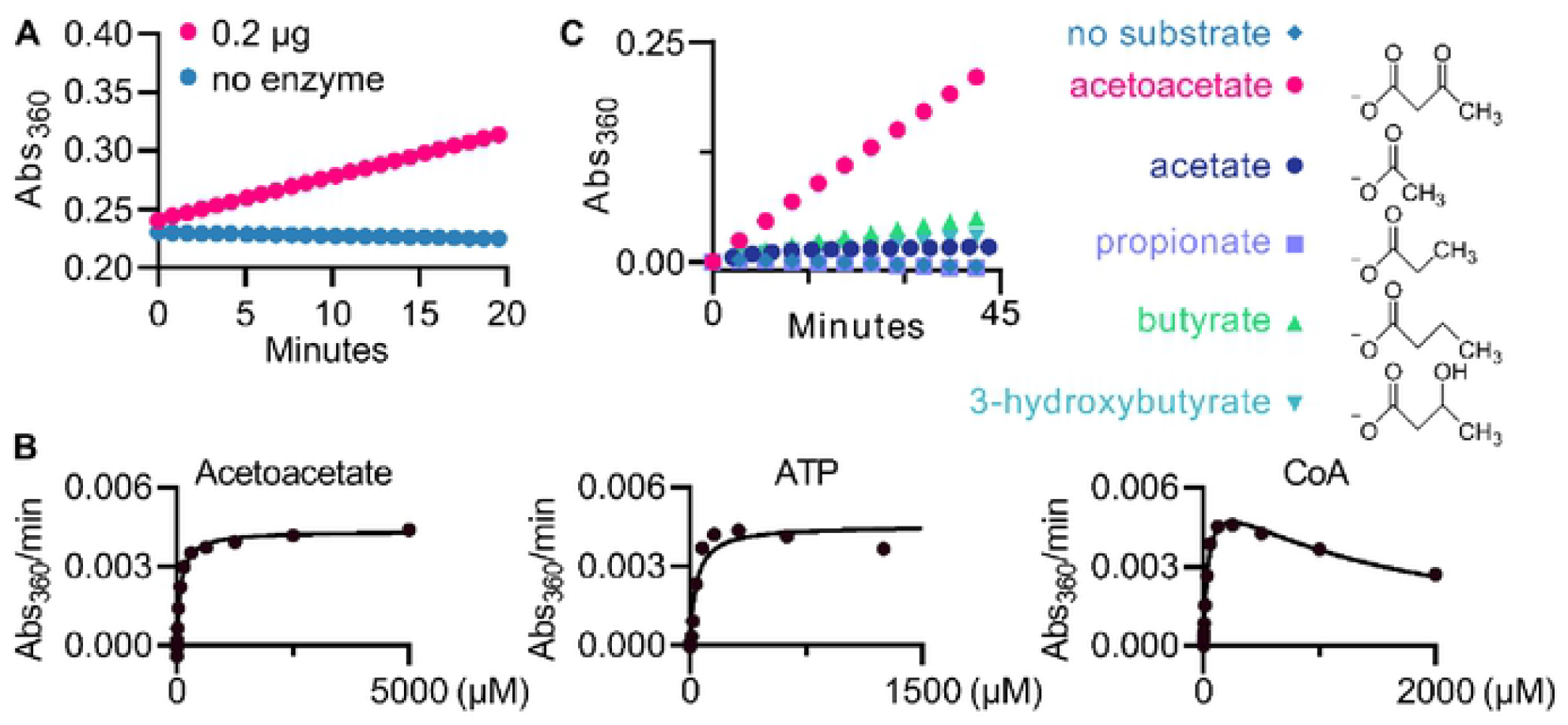
Biochemical characterization validates acetoacetate substrate preference of Kbc1. Reaction progression with 2 μg purified Kbc1 shows **a** long, steady linear range (A). Representative Michaelis-Menten curves for all substrates of Kbc1: acetoacetate, ATP, and CoA, respectively (B). Kbc1 activity using different substrate substitutions for acetoacetate; all substrates at 10 mM except for acetoacetate at 1 mM (C).

Despite the presence of the relatively small Gly residue in the putative substrate binding pocket, previously characterized acetoacetyl-CoA synthetases display selectivity for acetoacetate relative to other small-to-medium sized alkyl or aryl carboxylic acids (19). To assess the selectivity of CNAG_02045, we compared the activity of four carboxylic acids at 10 mM to acetoacetate at 1 mM (~5X K_m_). Acetate (C2), propionate (C3), butyrate (C4) and 3-hydroxybutyrate, the reduced form of acetoacetate and also a ketone body, showed minimal conversion under these conditions (Fig. 2C). CNAG_02054 is a selective acetoacetyl-CoA synthetase and is unlikely to function as a second acetyl-CoA synthetase. To align the gene name for CNAG_02045 with its biochemical function, we propose changing *ACS2* to *KBC1* which is based on the systematic IUPAC name for its substrate, 3-ketobutanoate, to both keep the three-letter convention and avoid confusion with acetyl-CoA synthetases.

### *ACS1* and *ACL1* appear to be synthetic lethal in *C. neoformans*

Acetyl-CoA is derived from multiple carbon sources (Fig. 3A). Glucose, the preferred carbon source for *C. neoformans* (13), can be converted to acetyl-CoA by two metabolic pathways (9, 12). In both pathways, glucose is glycolytically converted to pyruvate. In the first pathway, pyruvate is directly converted to acetyl-CoA in the mitochondria by pyruvate dehydrogenase and enters the tricarboxylic acid (TCA) cycle. Acetyl-CoA cannot cross cellular membranes (20) and, thus, the acetyl-CoA equivalent is exported from the mitochondria to the cytosol as citrate. *ACL1* then converts citrate to succinate and one molecule of acetyl-CoA (21). In the second pathway, pyruvate is converted to acetaldehyde by pyruvate decarboxylase. Aldehyde dehydrogenases oxidize acetaldehyde to acetate which is the substrate for *ACS1*.

**Figure 3:**
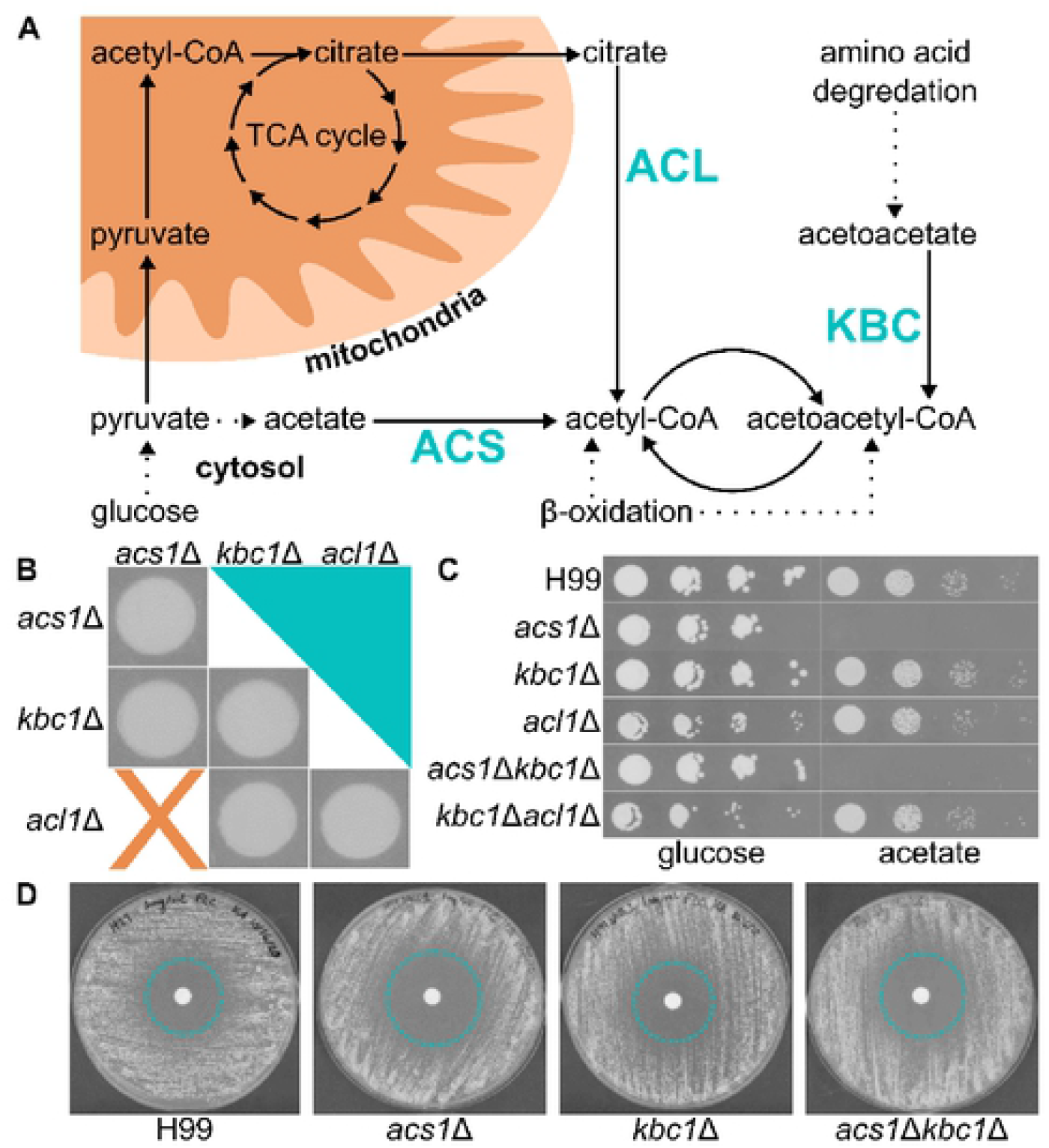
Acetyl-CoA production requires *ACL1* or *ACS1*, but not *KBC1* in vitro. Diagram of major acetyl-CoA sources in *C. neoformans* (A). Different single and double deletion mutants made, spots on YPD were imaged after 48 hours at 30°C (B). Spot dilutions of all mutants on YNB supplemented with either 2% of glucose or acetate as the carbon source, imaged after 72 hours at 30°C (C). Disk diffusion with 20 μg fluconazole on YNB +2% glucose plate, imaged after 72 hours at 37°C (D).

The oxidation of fatty acids generates one molecule of acetyl-CoA per round of β-oxidation and ultimately generates acetoacetyl-CoA as the final product. Acetoacetate is generated by the degradation of ketogenic amino acids such as leucine. *KBC1* converts acetoacetate to acetoacetyl-CoA and thus these two pathways converge. Acetoacetyl-CoA lyase (*POT1*) converts acetoacetyl-CoA to two molecules of acetyl-CoA. To explore how these pathways function together, we generated a systematic set of double deletion mutants derived from *ACL1, ACS1*, and *KBC1* (Fig. 3B). We were able to construct deletions for all combinations except *acl1*Δ *acs1*Δ. Despite multiple attempts including the generation of a conditional expression allele regulated by the *CTR4* promoter, we were unable to construct a strain lacking both *ACL1* and *ACS1*. This apparent synthetic lethality suggests that these two pathways compensate for one another and are the two dominant pathways under laboratory conditions.

We screened the single and double mutants on a wide range of media and growth conditions. We found no standard growth media or carbon sources in which the *kbc1*Δ alone or in combination led to a growth defect relative to WT or the single mutant of either *acl1*Δ or *acs1*Δ. Representative examples are shown in Fig. 3C and a complete list of conditions is provided in Table S1. This is likely because it is very difficult to establish conditions where the cell is completely dependent upon amino acid degradation for carbon. In addition, the degradation of leucine generates one molecule of acetyl-CoA during the conversion of HMG-CoA to acetoacetate and, thus, Kbc1 is not completely essential for the conversion of leucine to acetyl-CoA.

One of the critical functions of acetyl-CoA is as a precursor to ergosterol biosynthesis. We, therefore, determined the susceptibility of the double mutants to fluconazole. Interestingly, we did find that the *acs1*Δ mutant was modestly but consistently more susceptible to fluconazole than WT with a 2-fold reduction in minimum inhibitory concentration (MIC) under modified CLSI conditions (MIC: WT, 4 μg/mL; *acs1*Δ, 2 μg/mL). We confirmed this observation using disc diffusion assays (Fig. 3D). The deletion of *kbc1*Δ did not affect the MIC. Hu et al. previously reported that the *acs1*Δ mutant showed the same fluconazole susceptibility as WT (9). We suspect that the difference is that Hu et al. determined MICs at 30°C in YNB and YPD medium while we used buffered RPMI and incubated the cells at 37°C. Regardless, the loss of *ACS1* has a modest effect on fluconazole susceptibility in vitro.

### Deletion of *ACS1* and *KBC1* reduces survival in macrophages

Previously, Hu et al. reported that deletion of *ACS1* had no effect on the three classic *C. neoformans* virulence traits: growth at 37°C, melanin formation, or capsule formation (9). Similarly, deletion of *KBC1* either alone or in combination with the *acs1*Δ mutation did not affect these phenotypes. We did not examine the deletion of *KBC1* or *ACS1* in combination with *acl1*Δ because the *acl1*Δ mutant shows severe virulence trait phenotypes (12) and therefore it would not be possible to detect additional changes in the double mutants.

A critical aspect of *C. neoformans* pathogenesis is its ability to replicate within the macrophage (3). The macrophage is thought to be a nutrient-poor niche (22) and, therefore, we hypothesized that it may place demands on central carbon metabolism that are not well-replicated in vitro (23). We, therefore, tested the ability of the double mutants to replicate within the murine macrophage-like cell line J774.

All mutants, with the exception of the *acl1*Δ mutant, were phagocytosed by J774 cells similarly to the H99 parental strains (Fig. 4A). Griffths et al. had found that there was no difference in uptake of their *acl1*Δ mutants using identical media and opsonization. It is not clear what the distinction in our results may be. In contrast, the intracellular survival of the a*cs1*Δ, *kbc1*Δ, *acl1*Δ, *acs1*Δ *kbc1*Δ, and *acl1*Δ *kbc1*Δ mutants within J774 cells were reduced relative to H99 by ~2-3-fold (Fig. 4B). There were no significant differences between the different mutants. These data are consistent with the nutrient poor status of the macrophage environment. Stringency of the nutrient stress is such that loss of any of the three acetyl-CoA generating pathways leads to reduced fitness in this key *C. neoformans* niche.

**Figure 4:**
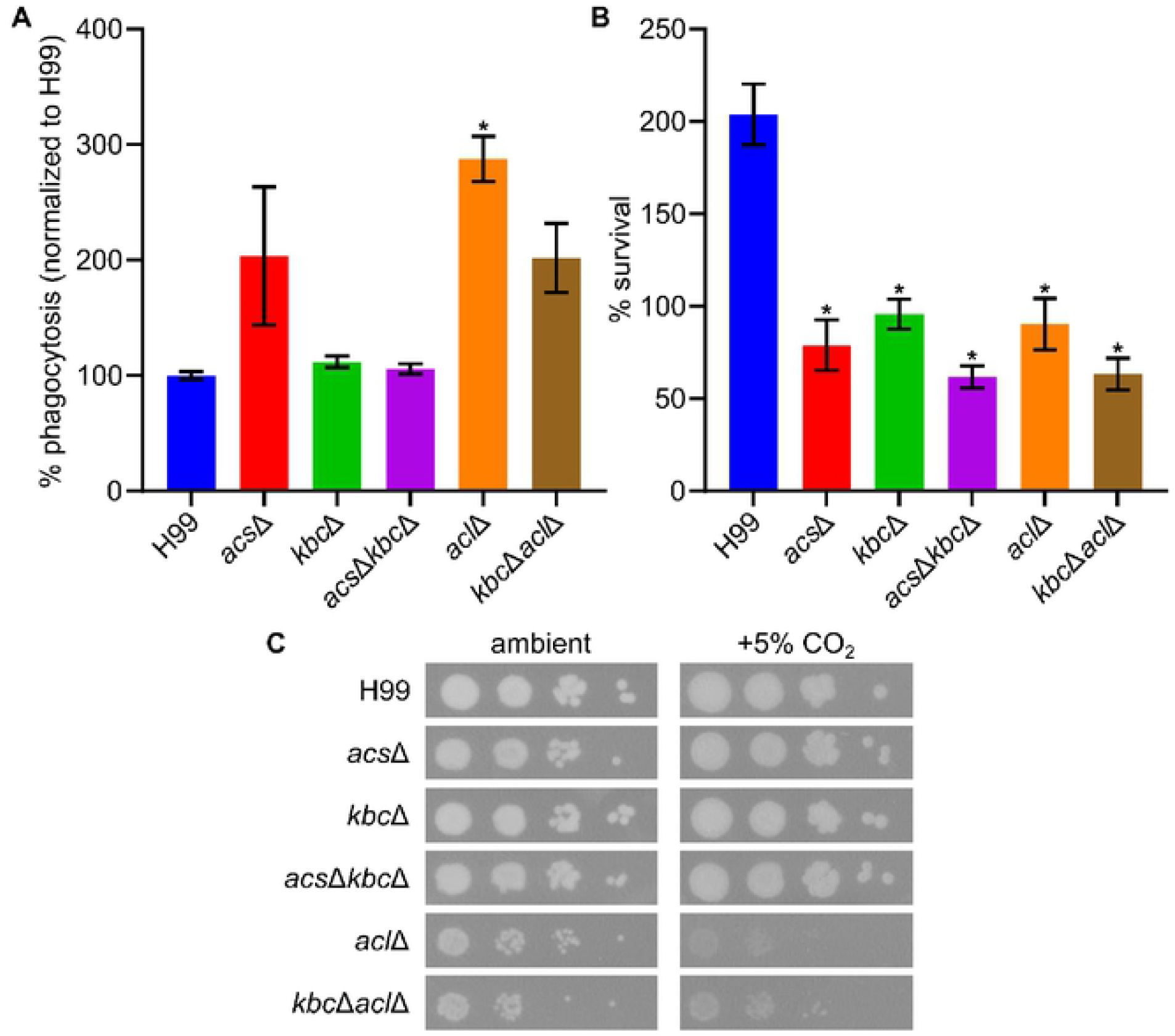
*ACL1, ACS1*, and *KBC1* are all required for full fitness in phagocytosed cells. Phagocytosis of different mutants by J774 (MOI 5:1) macrophages normalized to H99 parent strain (A). Survival of phagocytosed mutants in J774 cells 24 hours after uptake, normalized to each strain’s phagocytosis levels (B). Spot dilutions of *acl*Δ mutants show a growth defect on RPMI-MOPS with 5% CO_2_, plates imaged after 96 hours at 37°C ± 5% CO_2_ (C). Analysis done using a one-way ANOVA; asterisks (*) indicate P < 0.05 compared to H99.

To further explore the phenotypes of the acetyl-CoA-generating mutants under host-like in vitro conditions, we compared their growth on buffered RPMI media at 37°C under both ambient and host concentrations of carbon dioxide (5%). The *acl1*Δ mutant had a modest growth defect that was increased by the deletion of *KBC1* but not *ACS1* under ambient concentrations of carbon dioxide. The growth defects of the *acl1*Δ and *acl1*Δ *kbc1*Δ mutants were exacerbated significantly at host concentrations of carbon dioxide, indicating that *ACL1* is critical for the cells to tolerate the stress of elevated carbon dioxide in the host. These data also further support the notion that the importance of the three enzymes varies with the specific environmental conditions and stresses.

### *KBC1* and *ACS1* expression is increased during infection and host-like in vitro conditions

To further characterize the interplay between the different acetyl-CoA-generating enzymes, we examined their relative expression levels. As discussed above, *ACS1* has previously been shown to be expressed at much higher levels in the mouse lung than in standard laboratory culture conditions, while *ACL1* expression is relatively stable (10). More recently, *KBC1* was also found to be highly expressed in CSF both in vivo and ex vivo (11). These experiments used large-scale methods and, therefore, we were interested in confirming them and looking directly in infected brain tissue using RT-PCR. We also examined tissue at later time points than Hu et al. had used. Consistent with previous results, *ACS1* expression is much higher in infected lung (inoculated intra-tracheally) than in RPMI (Fig. 5A). Similarly, *KBC1* expression is also upregulated dramatically in the lung. *ACS1* is not significantly upregulated at the time point we examined in the brain after intravenous infection. However, *KBC1* expression in the brain is much higher than in RPMI culture. These data suggest that central carbon metabolism is significantly different during infection compared to in vitro conditions.

**Figure 5:**
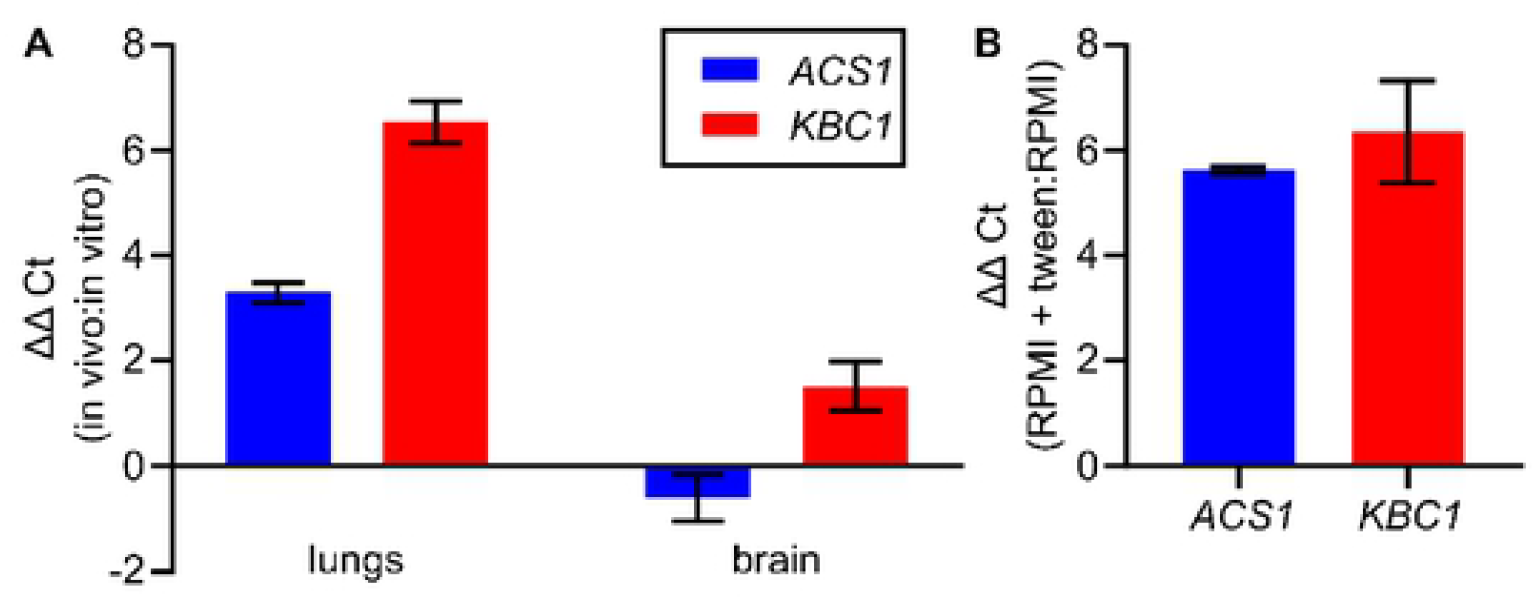
*ACS1* and *KBC1* expression is increased in vivo and in vitro in the presence of fatty acid. In vivo *ACS1* and *KBC1* expression from lungs recovered 17 days after intranasal inoculation and brains recovered 8 days after intravenous inoculation compared to in vitro cultures grown to early log phase in RPMI at 30°C (A). *ACS1* and *KBC1* expression is induced by the addition of a soluble lipid (tween-20) to RPMI (B).

Based on these findings, we examined a number of host-like in vitro conditions (e.g., RPMI, RPMI+5% CO_2_) to identify those that induce expression of *KBC1* and *ACS1* but were not initially successful at doing so. However, we reasoned that both the lung and brain have relatively high amounts of lipid. To test if lipids induce expression of *KBC1* and *ACS1*, we supplemented RPMI with tween-20 as a soluble source of fatty acid. As shown in Fig. 5B, this increased expression of both *KBC1* and *ACS1* relative to RPMI alone albeit not to the extent that we observed in vivo. This observation suggests that, in the presence of fatty acids, multiple pathways for acetyl-CoA generation are induced and that may contribute to the increased expression of *KBC1* and *ACS1* in vivo. We did not, however, observe a significant growth defect in the *kbc1*Δ, *acs1*Δ, or *kbc1*Δ *acs1*Δ double mutant in RPMI supplemented with tween-20 relative to RPMI alone; however, the lipids are not the only carbon source under these host-like conditions.

### Loss of both *ACS1* and *KBC1* reduces fitness during CNS infection and increases susceptibility to fluconazole

The increased expression of *ACS1* and *KBC1* during infection suggested that they may play a more important role in the fitness of *C. neoformans* in vivo as compared to in vitro. Furthermore, these mutants showed defects in fitness within macrophages. Since the *acs1*Δ mutant is modestly more susceptible to fluconazole, we were also interested to see if this phenotype was manifest during infection. Hu et al. had already shown that deletion of *ACS1* led to reduce virulence in a pulmonary infection model of cryptococcosis (9); deletion of *KBC1* in the *acs1*Δ background did not show any additional effects (data not shown). We, therefore, asked if the genes might interact during disseminated infection to the CNS. As shown in Fig. 6A, the fungal brain burden of mice infected with neither the *acs1*Δ nor *kbc1*Δ mutant differed from those infected with wild type. In the presence of fluconazole, however, the *acs1*Δ mutant was more susceptible to fluconazole in vivo than either WT or the *kbc1*Δ mutant by approximately 0.5 log_10_ CFU/mL.

**Figure 6:**
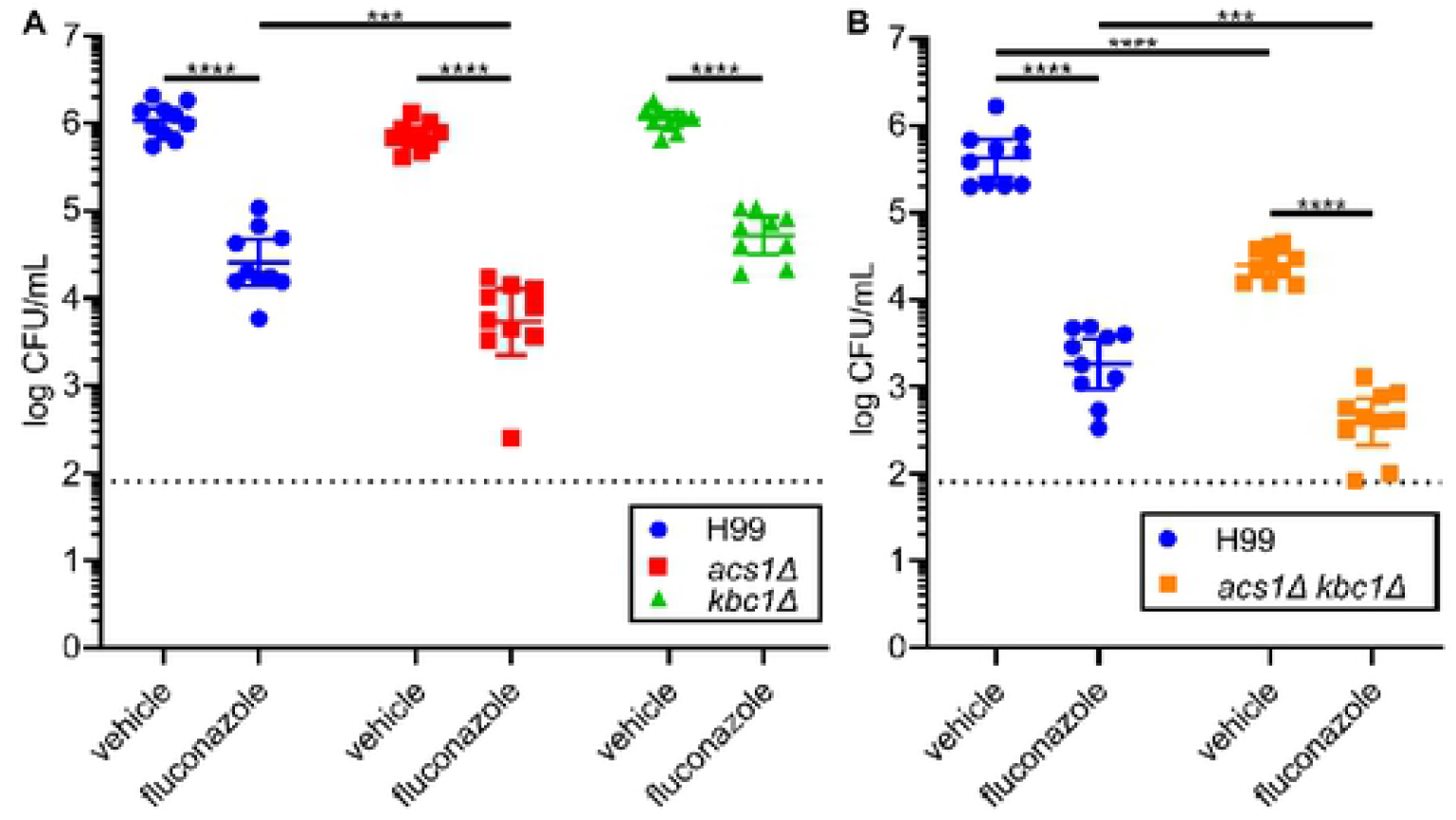
Alternative Pathways for acetyl-CoA generation are required for full fitness during brain infection and affect in vivo fluconazole susceptibility. CD-1 mice were inoculated with either the *acs1*Δ or *kbc1*Δ mutant and treated with either 125 mg/kg fluconazole or vehicle for 2 days when their brains were harvested, homogenized, plated, and growth at 30°C for 48 hours before counting (A). The experiment was repeated with the *acs1kbc1* double mutant (B). Analysis done using a two-way ANOVA; three asterisks (***) indicate 0.0001 < P < 0.001 and four asterisks (****) indicate P< 0.0001.

In contrast to the two single mutants (Fig. 6B), the fungal burden of the *acs1*Δ *kbc1*Δ double mutant was reduced by > 2 log_10_ CFU/mL, indicating that these two acetyl-CoA-generating pathways likely compensate for one another during infection of the CNS. In the presence of fluconazole, the fungal burden of the double mutant is further reduced to near the limit of detection of the infection assay. Taken together, these data clearly indicate that acetyl-CoA biosynthesis is dependent upon multiple pathways during CNS infection in a manner that could not be predicted from in vitro characterization. Furthermore, these data also indicate that the metabolic pathways required for pulmonary infection are not necessarily the same as those required for disseminated infection of the CNS.

## Discussion

Acetyl-CoA is one of the most fundamental molecules in biology, in general, and in carbon metabolism, in particular. Fungi can be distinguished into two groups based on the types of enzymes that generate acetyl-CoA (9, 12, 15). The first group relies on ACS localized to nucleus/cytoplasm and mitochondria to generate non-pyruvate decarboxylase-derive molecules acetyl-CoA. This group includes *Saccharomyces cerevisiae* and *Candida albicans*. In these yeast species, acetyl-CoA synthetases are essential (24). The second group of fungi generate acetyl-CoA through two enzymes: ACS and ACL. *C. neoformans* and *Aspergillus* spp. are two examples of fungi with both ACS and ACL (25). In humans, ACL is the major source of acetyl-CoA while ACS contributes significant acetyl-CoA only under severe stress or in the case of tumor cells where it can become the major acetyl-CoA generating enzyme (26). ACL is essential in mammals (27).

Our group has identified a molecule, AR-12, that inhibits fungal ACS and has potent, broad spectrum antifungal activity including fungi that have both ACS and ACL (14). In mouse model efficacy studies, AR-12 improves the activity of fluconazole against *C. neoformans*. Since ACS is typically not essential in fungi with both ACS and ACL enzymes, we took a genetic interaction approach to investigate the mechanistic basis for the antifungal activity of AR-12. We have been unable to generate double mutants of *ACS1* and *ACL1* or conditional expression strains of either gene in the absence of the other. Although not definitive, these observations strongly suggest that these two genes are synthetically lethal. As reported previously by Kronstad lab (12), mutants lacking *ACL1* have increased expression of *ACS1*, suggesting that these two enzymes may compensate for one another and further supporting the assertion that loss of both enzymes would be lethal.

Acetyl-CoA equivalents generated in the mitochondria by the TCA cycle are exported from to the cytosol as citrate which is then converted to acetyl-CoA and oxaloacetate by Acl1. As such, conditions that require Acl1-generated acetyl-CoA are also dependent upon mitochondrial function. In our characterization of the mode of action for AR-12, we discovered that AR-12 rapidly depolarizes the mitochondrial membrane (14), leading to a loss of the proton-motive force. The electron-transport chain is required to replenish NAD^+^/FAD^+^ for proper TCA function and, thereby, deliver citrate to the cytosol for conversion to acetyl-CoA by Acl1. Consequently, AR-12 may induce the equivalent of a reduction in Acl1 function by interfering with the ability of the cell to generate citrate through the TCA cycle. Therefore, we propose that the broad-spectrum activity of AR-12 may be due to the direct inhibition of Acs1 coupled with the indirect reduction in substrate for Acl1, leading to the equivalent of a synthetic lethal interaction.

Cytosolic acetyl-CoA is required for lipid biosynthesis, including ergosterol. Consistent with this requirement, Griffiths et al. found that *acl1*Δ deletion mutants were hypersusceptible to fluconazole (12). Under the conditions that they used (YPD and YNB at 30°C), *acs1*Δ mutants did not show significant changes in susceptibility; at 37°C on buffered RPMI medium, *acs1*Δ mutants showed modestly increased susceptibility to fluconazole. Deletion of *KBC1* in the *acs1*Δ background did not further sensitize the strain to fluconazole. One of the striking findings of our work is that the effect of these mutations on fluconazole susceptibility is much more pronounced in vivo than it is in vitro (Fig. 6). The *acs1*Δ mutant has only a 2-fold change in MIC in vitro but the brain burden of *acs1*Δ-infected mice is reduced by 5-fold. Again, in contrast to in vitro conditions, deletion of *KBC1* in the *acs1*Δ background further increased the fluconazole susceptibility to the point where the infection was nearly cleared. These data indicate that during brain infection ergosterol synthesis is likely to be dependent on all three acetyl-CoA-generating enzymes.

Further supporting this conclusion, the *kbc1*Δ *acs1*Δ double mutant has a fitness defect in the absence of antifungal drugs. Deletion of *ACL1* is already known to severely reduce fitness in vivo, indicating it plays a key role in the generation of acetyl-CoA in vivo. However, the fitness defect of the *kbc1*Δ *acs1*Δ strain indicates that Acl1 is not sufficient to be the sole source of acetyl-CoA during brain infection. Although the mechanistic basis for this observation remains unclear, our expression studies indicate that the expression of *KBC1* and *ACS1* is increased during infection and under host-like in vitro conditions supplemented with lipid. Taken together, these data suggest that *C. neoformans* may be generating acetyl-CoA from multiple pools of carbon during infection.

The macrophage is another key niche for *C. neoformans* and the ability to replicate within the phagolysosome is a critical part of the mechanism of pathogenesis (3, 22). The phagolysosome is widely regarded to be a nutrient-poor environment and phagocytosed yeast show transcriptional profiles that support this model (23). Again, Griffiths et al. found that genes required for acetyl-CoA synthesis were strongly induced (12), including *ACL1*. Consistent with these findings, they also found that the *acl1*Δ mutant showed reduced replication within macrophages. Although neither *ACS1* nor *KBC1* was induced in macrophages, both genes are also required for full fitness in macrophages. Based on the observation that loss of any of the three acetyl-CoA-generating enzymes reduces fitness, the macrophage environment appears to be the most stringent in terms of the number of pathways required to maintain sufficient acetyl-CoA production. The utilization of multiple carbon metabolism pathways during macrophage infection by C. neoformans is consistent with recent observations reported for C. albicans as well (22, 23).

The only in vitro genetic interaction that we observed was between *ACL1* and *KBC1* on RPMI medium at ambient carbon dioxide. The main carbon sources present in this medium are glucose (0.5%) and amino acids. *ACL1* deletion reduces growth in the presence of this low concentration of glucose on multiple media and we suspect that the further reduction in growth may be because Kbc1 is part of the pathway converting amino acids (e.g., leucine) to acetyl-CoA. It is also interesting that host levels of carbon dioxide further exacerbate the fitness defect of *acl1*Δ on host-like media. Our group has recently shown that host concentrations of carbon dioxide is an independent stress with relevance to virulence (8). Specifically, environmental strains with low mammalian virulence do not tolerate host concentrations of carbon dioxide while clinical strains do. The decreased fitness of the *acl1*Δ mutant indicates that carbon dioxide increases the cell’s dependence on acetyl-CoA derived from the mitochondria. Our observation of this phenotype provides additional insight into the basis for the profound virulence defect observed for strains lacking *ACL1*.

Price et al. previously showed that the glycolytic pathway was critical for disease but not persistence within the the CNS (13). In the absence of glycolysis, alternative pathways to the generation of acetyl-CoA would be required. Our data strongly suggest that a combination of Acs1 and Kbc1 provide this carbon flow and, thereby, support the replication of *C. neoformans* within the CNS. As such, our observations for the role of Acs1 and Kbc1 fit well with existing data and provide additional insights into the metabolic requirements of *C. neoformans* during infection.

In summary, our genetic interaction approach to the study of three enzymes involved in acetyl-CoA production indicate that *C. neoformans* uses a variety of carbon sources and pathways to generate this critical metabolite during infection. Although *C. neoformans* is well-known to have a strong preference for glucose utilization and is an obligate aerobe, it appears to also have significant metabolic versatility which, in turn, is particularly important during infection of macrophages and the central nervous system.

## Acknowledgements

The authors thank Xiaorong Lin for helpful discussion. This work was supported by NIH grants 5R01AI147541 (DJK), 1R01AI161973 (DJK), T32AI007511 (AJJ), and 5F32AI145160 (SRB). The authors thank the students of the Marine Biological Laboratory Molecular Mycology of 2015 for generating the initial isolates of the acs1Δ mutant and Virginia Glazier for confirming the strains. The funders had no role in study design, data collection and interpretation, or the decision to submit the work for publication.

## Methods

### Strains, Media, and Growth

Strains from *Cryptococcus neoformans* H99 background were maintained in glycerol stocks kept at −80°C and recovered on YPD plates (1% yeast extract, 2% peptone, 2% dextrose, 2% agar). Experiments performed with the strains (with the exception of in vivo experiments, which were grown for ~48 hours) used cultures grown overnight shaking at 220 rpm at 30°C in 2 mL liquid YPD (1% yeast extract, 2% peptone, 2% dextrose). For growth on plates, YNB plates (0.17% yeast nitrogen base without amino acids, 0.5% ammonium sulfate, 2% agar) were supplemented with 2% m/v of a carbon source: glycerol, acetate, glucose, or BHB; overnights of desired strains were then diluted in liquid YPD to an OD_600_ of 1.0 and serial diluted 10-fold down to 0.001 OD_600_, all four dilutions of each strain were plated using a replica spotter onto the medium of choice. Incubations under a carbon dioxide atmosphere (5%) were carried out in a standard tissue culture incubator at 37°C.

### Strain construction

*ACS1* deletions were performed by replacing the gene locus with the nourseothricin resistance marker using biolistics as previously described (28). To generate *KBC1* deletions in H99 and Δ*acs1*, both strains were transformed using the protocol and CRISPR-TRACE system previously described, with minor modification (29). The neomycin selective resistance marker was used to replace the *KBC1* locus using electroporation with a BioRad MicroPulser on the “Pic” setting. Strains were passaged three times on non-selective media before replating on selective media to check for stable marker integration. Selected mutants from both backgrounds were confirmed by southern blot. *ACL1* deletions were generated in the H99 and Δ*kbc1* strains using the *Cryptococcus* optimized Cas9 system with 50 base pair homology as previously described using a hygromycin resistance marker (30). The protocol for transformation was the same as before and selected mutants were again confirmed via southern blot.

### Enzyme expression and purification

*Cryptococcus neoformans KBC1* was cloned into the NdeI/XhoI site of the pET15-b expression vector and transformed into *Escherichia coli* strain BL21 with ampicillin selection. An overnight starter culture was prepared in LB broth with antibiotic and grown at 37°C at 220 rpm overnight. The next day, the overnight was diluted 1:100 in LB medium with antibiotic and grown to OD_600_ of 0.5-0.8, then induced with 1 mM isopropyl-β-D thiogalactopyranoside (IPTG) for two hours. Pelleted cells were resuspended in lysis buffer (1 mM PMSF, 1 mM DTT, 0.25 μL/L benzonase, 1 mg/mL lysozyme, 10 mM tris-HCl pH 7.5, 20 mM imidazole, 1 mM MgCl_2_, 200 mM NaCl) on ice and sonicated for two cycles of three minutes on, three minutes rest. Pelleted lysate was then mixed with washed Ni beads and transferred to a column. Wash buffer (20 mM imidazole, 20 mM tris-HCl pH 7.5, 150 mM NaCl) was run over the column, followed by elution buffer (300 mM imidazole, 20 mM tris-HCl pH 7.5, 150 mM NaCl). The elution was collected and placed in dialysis cartridges in Kbc1 enzyme buffer (200 mM NaCl, 50 mM tris-HCl pH 7.5, 1 mM MgCl_2_, 10% glycerol, 1 mM DTT) stirring overnight at 4°C. Purified Kbc1 was stored at −80°C.

### Acetoacetyl-CoA synthetase assay

Kbc1 activity was detected using our assay previously described for Acs1 activity except sodium acetoacetate was used in place of sodium acetate (18). Michaelis-Menten constants were determined for the acetoacetate, CoA, and ATP substrates by first determining the optimal concentrations of these substrates so that they would be in excess without, in the case of CoA, being high enough to inhibit the reaction. Substrates provided in excess allow the apparent Km to closely approximate the actual Km. The EnzChek Pyrophosphate Assay Kit (Thermo) was used with reagents prepared by manufacturer’s standards, and with 4 mM MgCl_2_, 10 mM DTT, 4.5 μg/mL CnKBC1, 100 μM CoA, and 200 μM ATP per 50μL reaction. The reagents were mixed and aliquoted at room temperature, allowed to incubate for 15 minutes at 37°C to mop background phosphate contamination, then acetoacetate (final concentration of 1 mM) was added to the reaction, and the plate was read continuously for 40 minutes at 37°C in a SpectraMax i3X Multi-Mode plate reader (Molecular Devices) at absorbance 360 nm. To test possible substrates, a dilution series of propionate, butyrate, 3-hydroxybutyrate, or acetate was added in place of acetoacetate.

### Assays of capsule formation and melanin production

H99, Δ*acs1*, Δ*kbc1*, and Δ*acs1*Δ*kbc1* strains were grown overnight in YPD. The next day, 1mL of each culture was spun down, washed twice in PBS, resuspended in 1mL of RPMI/MOPS (pH 7.4), counted using a digital hemocytometer (Countess), and diluted to 7.5 ×10^5^ cells per mL RPMI/MOPS. Diluted cells were then aliquoted 100μL per well of a 96 well plate and sealed with a permeable membrane and incubated at 37°C in 5% CO_2_. Time points were taken at 24, 48, and 72 hours, one well of each strain was recovered, spun down, supernatant removed, resuspended in 20 μL PBS, mixed 1:1 with India ink and imaged at 40x for capsule formation. For melanin production, overnights were diluted 1:100 in dH_2_O, counted using a Countess II FL (Invitrogen), and diluted to 1× 10^7^ cells per mL in water. A serial dilution in water was made down to 1× 10^3^ cells per mL and spotted onto L-DOPA plates using a replica spotter. Plates were incubated at 30°C for 2 days and then imaged.

### Antifungal drug susceptibility assays

Minimum inhibitory concentration assays were performed using a slightly modified CLSI standard method. Fluconazole was diluted 2-fold in DMSO and added to RPMI/MOPS so that the final DMSO concentration in each well was 1.28%. Overnights of selected strains were washed 3 times in PBS, resuspended in RPMI/MOPS, counted using a Countess II FL (Invitrogen), and diluted to 1 × 10^5^ cells per mL in RPMI/MOPS. Diluted cells were added to triplicate wells for each drug dilution such that each well had ~1000 cells and incubated at 37°C for 72 hours before checking for growth via visual inspection. Disk diffusion assays were done by taking 1 mL of overnights, washing 3 times in PBS, resuspending in 1mL PBS, then diluting 1:100 in PBS. The 1:100 dilution was spread on top of a YPD plate using a sterile swab to achieve a lawn of cells on the entirety of the plate. Sterile filter paper dots were then added on top of the lawns and 20 μL of desired drug or control was carefully pipetted on the dot and allowed to sink in. Plates were then incubated at 37°C for 72 hours and imaged.

### Gene expression analysis by quantitative reverse transcription PCR

For in vitro expression, 1 mL samples were taken in log phase, OD_600_ = ~0.3-0.65, from liquid cultures (with the exception of RPMI/MOPS + Tween-20 for H99, H99Δ*acs*, H99Δ*kbc*, and H99Δ*acs*Δ*kbc*, which were taken at stationary phase). With the MasterPure Yeast RNA Purification Kit (catalog number MPY03100), RNA was extracted using the manufacturer’s protocol. Pelleted cells were vortexed, mixed with 300 μL extraction reagent with 1 μL proteinase K, incubated at 70°C for 12 min with vortexing every 4 min, then placed on ice for 3-4 min. Samples were mixed with 175 μL MPC protein precipitation reagent and then were centrifuged for 10 min at 4°C and 12,000 rpm. Supernatants were transferred and mixed with 500 μL isopropanol, spun down again, rinsed with 70% ethanol twice and RNA was suspended in 35 μL TE buffer. Concentration was measured at 260 nm, and RNA was then converted to cDNA using iScript cDNA Synthesis Kit (Bio-Rad catalog #170-8891), with 4 μL 5x iScript reaction mix, 1 μL reverse transcriptase, 14 μL water, and 1-10 μL of RNA (so that there was ~0.4-1 μg RNA) per reaction.

For in vivo expression, A/J mice, 3 per condition, were inoculated with H99 1 × 10^4^ intravenously with brains harvested after 8 days or inoculated 5 × 10^4^ intranasally with lungs harvested after 17 days. Organs were lyophilized overnight and then bead beaten with zircon/silica beads for 45 seconds. 1 mL triazole reagent was added and tubes were incubated at room temperature for 10 min then spun at 10k rpm for 5 min at 4°C. The clarified layer was transferred to a new tube where 200 μL chloroform was added and samples were incubated for 5 min at room temperature then spun at 12k rpm for 15 min at 4°C. The top layer was transferred to a gDNA removal column from Qiagen RNeasy plus kit (Qiagen, catalog #74134) and the manufacturer’s instructions were then followed from that point. Samples were eluted in 50 μL water, diluted 1:10, and quantified. ~500 ng RNA was used to make cDNA with the same iScript cDNA synthesis kit; the cDNA was then diluted 1:1 in water.

Quantitative reverse transcription-PCR was done with cDNA using TEF1 as a housekeeping gene, *ACS1, KBC1*, and *ACL1* using IQ Sybr green supermix (Bio-Rad catalog #170-8882). Each sample & condition was in technical triplicate and biological duplicate, with 10 μL of the Syber green supermix, 0.25 μM of each respective primer, 7.5 ng/μL of cDNA from the in vitro samples or 2 μL of the 1:1 dilution from the in vivo cDNA, and 8 (in vitro) or 7 (in vivo) μL of water was in each reaction well.

### *C. neoformans-macrophage* interactions

J774 cells were seeded onto a 96-well plate at 2.25 × 10^5^ cells per well and incubated at 37°C with 5% CO_2_ for 24-48 hours until 80-90% confluent. Once they are confluent, standard overnight cultures of the desired yeast strains were washed twice with PBS, counted with the Countess II FL (Invitrogen), and diluted to 1.12 × 10^7^ CFU/mL in PBS for an MOI of 5:1. Each strain is then opsonized with antibody 18b7 (2 μg/mL final) and incubated at 37°C for 1 hour with rotation. After opsonization, 80 μL of cells per well were added to the two prepared J744 plates, each with 6 technical replicates. Plates were incubated for 80 min and then washed to remove cells that had not been endocytosed. To determine phagocytosis from the first plate, macrophage cells were lysed with 0.1% cold triton X for 10 min while rocking before plating. To then determine how well those that had been phagocytosed survived, J774 growth media was added to the second washed plate and left for 24 hours at 37°C with 5% CO_2_ and then lysed with 0.1% cold triton X for 10 min while rocking before plating. For plating, cells were serially diluted 10-fold down to 1:1000. All four dilutions (full down to 1:1000) for each well were plated on YPD with 3x 10 μL spots per dilution and grown for 72 hours at 30°C before counting CFU.

### Mouse infection model of cryptococcosis

All animal experiments were carried out in compliance with approval from the The University of Iowa Institutional Animal Care and Use Committee (IACUC). For fluconazole susceptibility, 50 mL cultures of strains were grown for 48 hours (YPD, 30°C, shaking) and then used to inoculate female CD-1 mice via tail vein injection with 200 μL containing 3.8 × 10^5^ cells. 20 mice were inoculated for with each strain, 10 to be treated with 125 mg/kg fluconazole, 10 to receive the vehicle. Mice were monitored and treatments of fluconazole or vehicle were given via intraperitoneal injection every 24 hours starting about 1 hour after the initial inoculation. After 3 treatments, the mice were euthanized with CO_2_ and their brains homogenized in 1mL PBS and serially diluted 10-fold down to 1:10,000. Dilutions of 1:10 down to 1:10,000 for each organ were plated on YPD with 6x 10 μL spots per dilution and plates were grown at 30°C for 48 hours and then CFU were counted.

